# Distinct encoding of decision confidence in human medial prefrontal cortex

**DOI:** 10.1101/251330

**Authors:** Dan Bang, Stephen M. Fleming

## Abstract

Our confidence in a choice and the evidence pertaining to a choice appear to be inseparable. An emerging computational consensus is, however, that the brain should maintain separate estimates of these quantities for adaptive behavioural control. Here we devised a psychophysical task to decouple confidence in a perceptual decision from both the reliability of sensory evidence and the relation of such evidence with respect to a choice boundary. Using human fMRI, we found that an area in medial prefrontal cortex (perigenual anterior cingulate cortex, pgACC) tracked expected performance, an aggregate signature of decision confidence, whereas neural areas previously proposed to encode decision confidence instead tracked sensory reliability (posterior parietal cortex and ventral striatum) or boundary distance (pre-supplementary motor area). Supporting that information encoded by pgACC is central to a subjective sense of decision confidence, we show that pgACC activity does not simply co-vary with expected performance but is also linked to within-subject and between-subject variation in explicit confidence estimates. Our study is consistent with the proposal that the brain maintains choice-dependent and choice-independent estimates of certainty, and sheds light on why dysfunctional confidence often emerges following prefrontal lesions and/or degeneration.

**Significance Statement:** Recent computational models propose that our sense of confidence in a choice reflects an estimate of the probability that the choice is correct. However, it has proven difficult to experimentally separate decision confidence from its component parts, such as our certainty about perceptual evidence or choice requirements. Here we devised a task to dissociate these quantities and isolate a distinct encoding of decision confidence in the medial prefrontal cortex of the human brain. We show that activity in this area not only tracks expected performance on a task but is also related to both within-subject and between-subject variation in a subjective sense of confidence. Our study illuminates why dysfunctional confidence often emerges following damage to prefrontal cortex.

## Introduction

Decisions are often made in the face of uncertainty and in the absence of immediate feedback. Accompanying such decisions is a sense of confidence in having made the right choice which can be used to guide behaviour (1, 2). For example, after having made a difficult choice, an animal might correctly estimate that its decision is unlikely to be correct and thus avoid wasting time waiting for a reward that may not arrive (3). Humans may communicate such estimates of decision confidence to make more accurate decisions together than alone (4). Despite widespread agreement that decision confidence is a useful quantity for adaptive control of behaviour, neurobiological support for a distinct encoding of decision confidence is lacking.

Several computational models propose that decision confidence reflects an internal estimate of the probability that a choice is correct (1, 2). A ubiquitous paradigm for studying the neural basis of this computation is sensory psychophysics. On a typical trial, subjects first make a categorical choice about an ambiguous stimulus, such as deciding whether a cloud of dots is moving left or right or whether a contrast grating is tilted counterclockwise or clockwise of vertical. Subjects then make a secondary judgement which requires estimating the probability that the initial choice is correct. For example, subjects may have to decide whether to opt out of the choice for a sure but small reward or, in humans, explicitly estimate their confidence in the choice. Variation in these confidence-based behaviours may be induced by manipulation of task features, such as stimulus reliability (e.g., percentage of coherently moving dots) or the distance between the stimulus and a choice boundary (e.g., angular deviation from vertical axis), and/or intrinsic stochasticity in neural processing.

This general approach has been used with various species and techniques to identify neural areas which are involved in, or at least predict, confidence-based behaviours. In monkeys, single-unit recording and functional inactivation have identified lateral intraparietal sulcus (5), thalamic pulvinar (6), supplementary eye fields (7) and dopaminergic neurons in substantia nigra (8) as contributing to confidence-based behaviours, such as opt-out responses and post-decision wagers. In rodents, similar approaches have identified orbitofrontal cortex as involved in confidence-based behaviours, such as willingness to wait for a potential reward (3). In humans, functional magnetic resonance imagining (fMRI) has identified neural areas which track explicit confidence estimates, including striatum (9), medial prefrontal cortex (10, 11), dorsal anterior cingulate cortex (12) and rostrolateral prefrontal cortex (11, 12).

In the tasks discussed above, there are at least two distinct components to a computation of decision confidence: the reliability of sensory evidence, and the relation of such evidence with respect to a choice boundary (1). When the choice boundary is known in advance or fixed, one of these components may be a sufficient statistic for estimating decision confidence. For example, it has been proposed that decision confidence, algorithmically, is a function of the sensory evidence in favour of a choice and elapsed time (5) or the (absolute) distance between such evidence and a choice boundary (3, 6, 9). Such close coupling between decision confidence and its component parts can, however, make it hard to evaluate the contribution of distinct neural signals to confidence-based behaviours. For example, neural activity in parietal cortex might predict an animal’s opt-out responses (5) because the area encodes a probability distribution over sensory states given sensory evidence – a key component of decision confidence – rather than decision confidence per se. Conversely, neural activity in orbitofrontal cortex might predict an animal’s willingness to wait for a potential reward (3) because the area encodes the distance between sensory evidence and a choice boundary – another component of decision confidence – rather than decision confidence per se. While the distinction between these encoding schemes may sometimes prove moot, there are many situations in which decision confidence cannot be readily estimated from the choice process and/or must be represented separately (1), such as when a choice boundary is not known at the start of a trial (13) or when performing a series of interdependent choices (14).

Here we tested for a distinct encoding of decision confidence in the human brain by devising a task which isolates decision confidence from its component parts. Subjects performed a continuous-direction, variable-reference random dot-motion task which, on aggregate, separated the probability that a motion discrimination judgement was correct (expected performance) from the reliability of a percept of motion direction (sensory reliability) and the distance between a motion percept and a choice boundary (boundary distance). Our approach builds on previous behavioural paradigms which have examined components of decision confidence (e.g., stimulus mean and variance) (15, 16). We show, using fMRI, that both aggregate and single-trial signatures of decision confidence are tracked by medial prefrontal cortex (perigenual anterior cingulate cortex, pgACC).

## Results

### Experimental isolation of decision confidence

Subjects (*N* = 32) viewed a field of moving dots inside a circular aperture (Fig. 1). On each update of the motion display, a fraction of dots moved coherently in a pre-specified direction, sampled anew on each trial from the range 1-360 degrees, whereas the rest moved randomly. After the motion display, a line transecting the aperture appeared. Subjects had to decide whether the net direction of dot motion was counterclockwise (CCW) or clockwise (CW) to this reference. Finally, subjects indicated their confidence in the choice being correct.

**Figure 1.**
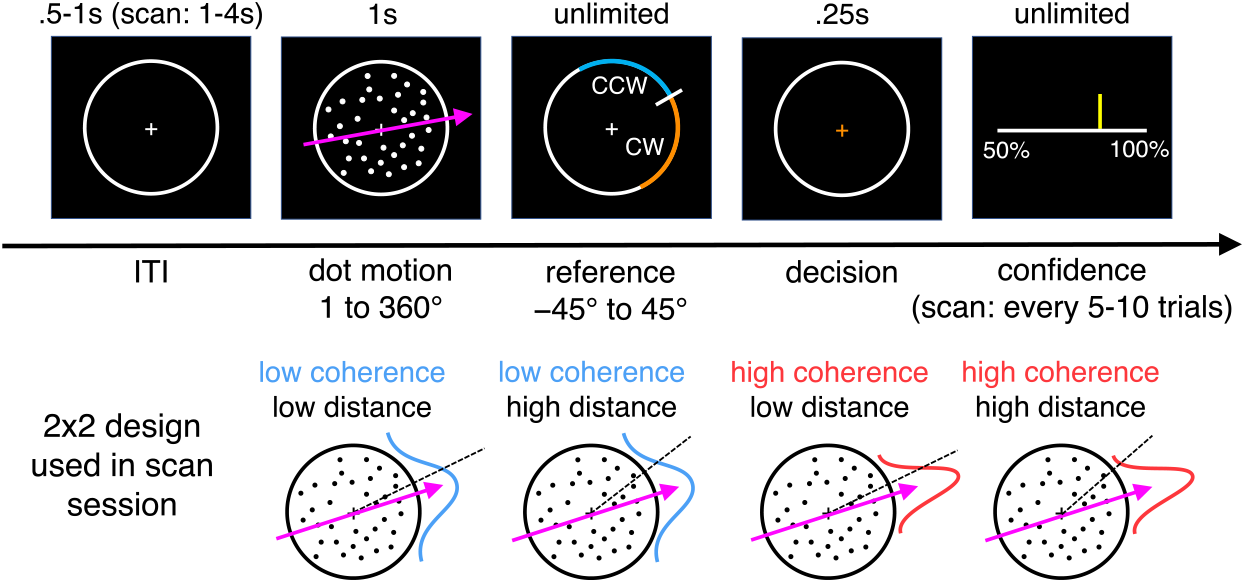
Continuous-direction random-dot motion task with variable reference. Subjects had to judge whether the net direction of dot motion was counterclockwise or clockwise to a reference which appeared after stimulus offset. We varied the percentage of coherently moving dots and the absolute angular distance between the motion direction and the reference. In the pre-scan session, confidence estimates (50% to 100% in steps of 10%) were elicited on every trial. In the scan session, confidence estimates were elicited every 5-10 trials. See **SI Appendix: Fig. S1** for information about stimulus calibration.

We varied, using a factorial design, the fraction of coherently moving dots (coherence) and the absolute angular distance between the motion direction and the reference (distance). Our aim was to separate a subject’s internal estimate of the probability that a motion discrimination judgement is correct from the reliability of their percept of motion direction and the distance between their motion percept and a choice boundary – components that bear on a confidence computation but are in and of themselves not sufficient for confidence estimation in our task.

Subjects performed the task in separate pre-scan and scan sessions. In the pre-scan session, we calibrated a set of coherences and distances (2×4 design) associated with target levels of choice accuracy and evaluated the behavioural effects of our task manipulation. In the scan session, we simplified the design to ensure sufficient trials per condition for fMRI analysis, using a subset of the calibrated task parameters (2×2 design). To avoid confounding neural responses related to decision confidence with neural responses related to explicit reports, we elicited confidence estimates every 5-10 trials.

### Behavioural validation of experimental approach

We first validated that subjects’ expected performance varied with changes in coherence and distance. Indeed, we found that choice accuracy was affected by both factors: subjects were more likely to be correct when coherence was high and when distance was high (**Fig. 2a**, logistic regression, coherence: *t*(31) = 11.46, *p* < .001, distance: *t*(31) = 19.00, *p* < .001, interaction: *t*(31) = 9.85, *p* < .001). These effects were mirrored in choice reaction time: subjects made faster decisions when coherence was high and when distance was high (Fig. 2b, linear regression, coherence: *t*(31) = −4.97, *p* < .001, distance: *t*(31) = −11.58, *p* < .001, interaction: *t*(31) = −9.26, *p* < .001). Critically, consistent with the aim of our design, the effects of coherence and distance on choice accuracy were reflected in explicit confidence estimates: subjects reported higher confidence when coherence was high and when distance was high (**Fig. 2c**, ordinal regression, coherence: *t*(31) = 8.45, *p* < .001; distance: *t*(31) = 11.15, *p* < .001; interaction: *t*(31) = 9.40, *p* < .001). These confidence effects survived controlling for choice reaction time and nuisance factors such as the initial position of the confidence marker and the cardinality of motion direction (**SI Appendix: Fig. S2**). Finally, the effects of coherence and distance on choice behaviour and confidence estimates were replicated in the scan session (**SI Appendix: Fig. S3**).

**Figure 2.**
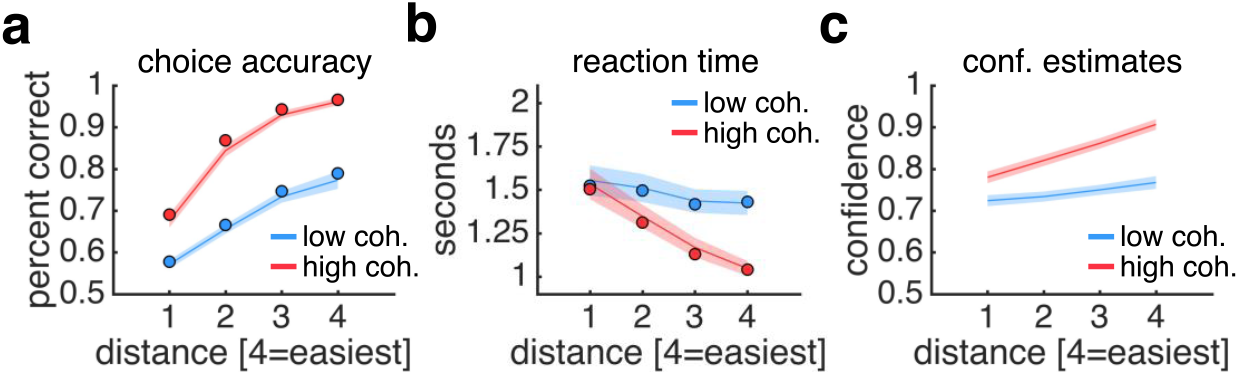
Behavioural results. **a**, Choice accuracy. **b**, Reaction time measured from reference onset. **c**, Confidence estimates. **a-b**, Solid dots are posterior predictives from a hierarchical drift-diffusion model fit to subjects’ responses separately for each condition. **a-c**, Data are from the pre-scan session. See **SI Appendix: Fig. S3** for equivalent plots from the scan session. Data are represented as group mean ± SEM.

To further unpack the drivers of the choice process, we modelled subjects’ responses using a hierarchical instantiation of the drift-diffusion model (DDM) (17). We remained agnostic as to how our task affected DDM parameters by fitting drift-rate (signal-to-noise ratio of evidence accumulation), threshold (amount of evidence needed for a choice) and non-decision time (e.g., stimulus encoding and response preparation) separately for each condition. In both sessions, we found that only drift-rate varied between conditions, whereas threshold and non-decision time were stable (see model predictions in **Fig. 2a-b** and parameter estimates in **SI Appendix: Fig. S4**). In our task, the momentary evidence entering into the accumulation process can be thought of as the signed difference between a noisy sample from a sensory representation of motion direction held in visual short-term memory and a choice boundary (18). Under this account, the reliability of the sensory representation, controlled by coherence, and the placement of the choice boundary, controlled by distance, *jointly* determine the signal-to-noise ratio of the accumulation process and thereby the probability that a choice is correct. Taken together, the DDM analysis shows that subjects’ choice strategy (threshold) and task engagement (non-decision time) were stable across conditions and sessions and supports that our task separates expected performance from sensory reliability and boundary distance.

### Isolating neural signatures of expected performance

Having validated our experimental approach, we next estimated a general linear model of the fMRI data. As shown above, the probability that a subject’s motion discrimination judgement is correct is a function of both the reliability of a subject’s percept of motion direction (coherence) and the distance between a subject’s motion percept and a choice boundary (distance) (**Fig. 2a**). We would therefore expect a brain region involved in tracking expected performance, an aggregate signature of decision confidence, to carry main effects of coherence and distance and, importantly, an interaction between these two factors. To identify such activity patterns, we adopted a masking approach. At a whole-brain level, we first searched for main effects of coherence and distance, and then, applying an *inclusive* mask constructed from the intersection of the two main effects (each map thresholded at *p* < .05, uncorrected), searched for a critical interaction between coherence and distance (*p* < .05, FWE-corrected). This analysis identified a single cluster in medial prefrontal cortex (perigenual anterior cingulate cortex, pgACC): in this area, activity tracked changes in both coherence and distance and, importantly, an interaction between these two factors (**Fig. 3a-b**), reflecting the pattern of both choice accuracy (**Fig. 2a**) and explicit confidence estimates (**Fig. 2c**)

**Figure 3.**
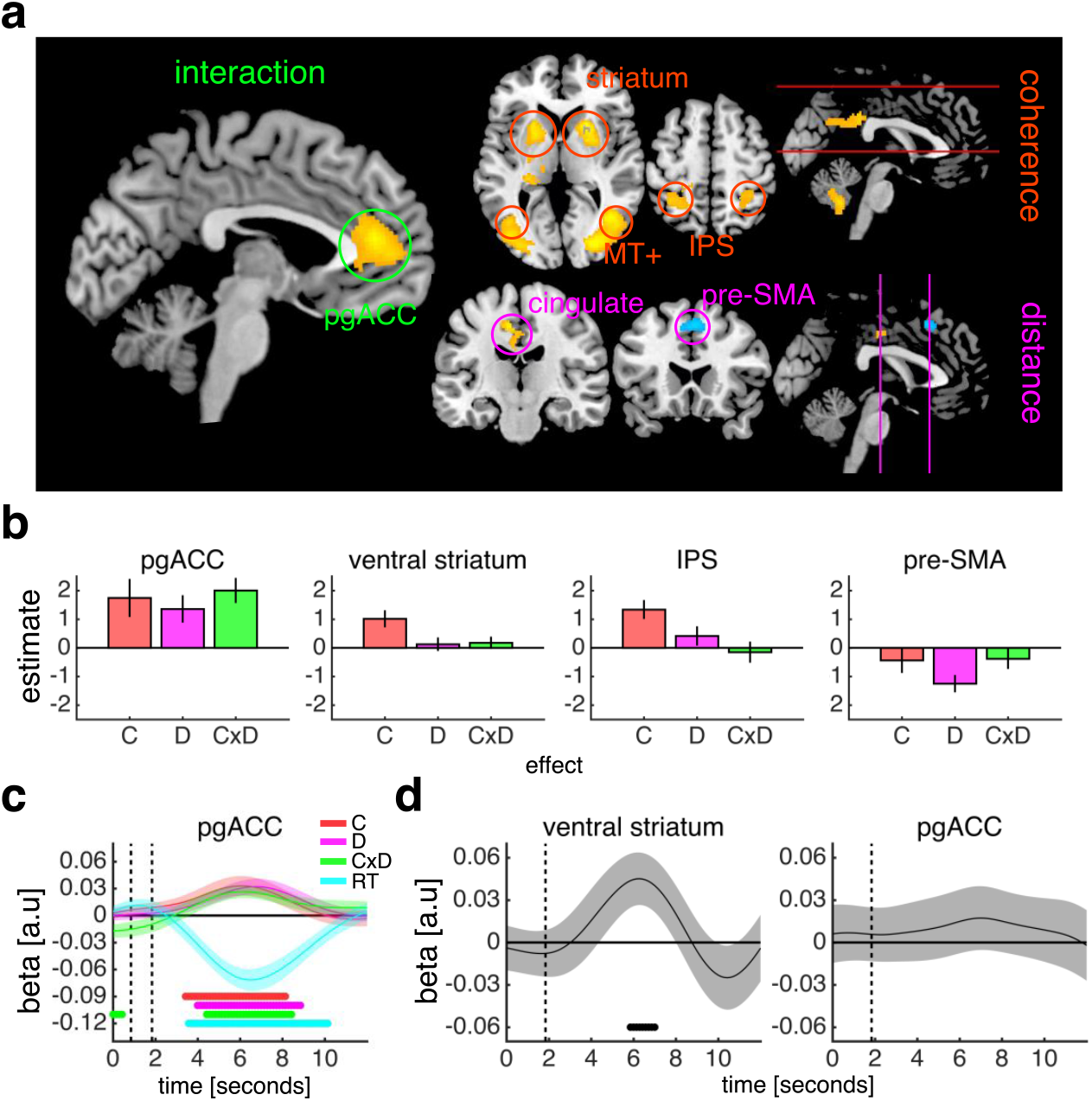
Neural signatures of expected performance, sensory reliability and boundary distance. **a**, Whole-brain factorial analysis of the effects of coherence, distance and the coherence × distance interaction. Activations are masked as detailed in the main text. Cluster colours denote positive (warm) and negative (cold) effects. Clusters are significant at *p* < .05, FWE-corrected for multiple comparisons; cluster-defining threshold: *p* < .001, uncorrected. Images are shown at *p* < .001, uncorrected. All clusters surviving whole-brain correction post- and pre-masking are detailed in **SI Appendix: Tables S1-2**. See **SI Appendix: Fig. S5** for control GLMs. **b**, ROI contrast estimates from factorial analysis of the effects of coherence (C), distance (D) and the coherence × distance interaction (C×D). c, General linear model analysis of the effects of coherence (C), distance (D), the coherence × distance interaction (CxD) and choice reaction time (RT) on ROI activity time courses. Vertical dashed lines indicate the onsets of the motion stimulus and the choice phase. See **SI Appendix: Fig. S6** for additional ROIs. d, General linear model analysis of the effect of reward magnitude on ROI activity time courses on confidence trials. Vertical dashed line indicates the onset of the reward magnitude cue. See **SI Appendix: Fig. S7** for additional ROIs. **b-d**, To avoid biasing subsequent analyses, ROIs were specified using simple contrasts from our factorial analysis (coherence, distance, coherence × distance) prior to masking, except for ventral striatum, which was specified anatomically. To avoid circularity, a leave-one-out cross-validation procedure was used for ROI specification. Data are represented as group mean ± SEM. **c-d**, Dots below time course indicate significant excursion of *t*-statistics assessed using two-tailed permutation tests.

We next identified areas which selectively tracked changes in coherence independently of distance. At a whole-brain level, we applied an *exclusive* mask constructed from the intersection of the main effect of distance and the coherence × distance interaction (each map thresholded at *p* < .05, uncorrected) and searched for a main effect of coherence (*p* < .05, FWE-corrected). This analysis identified clusters in extrastriate cortex, posterior cingulate cortex, parietal cortex and striatum, extending into the thalamus: in these areas, activity was higher when coherence was high but was unaffected by distance (**Fig. 3a-b**). The extrastriate and parietal clusters encompassed area MT+ and the intraparietal sulcus, respectively; areas which are sensitive to motion coherence and motion direction (19, 20).

Finally, we identified areas which selectively tracked distance independently of coherence. At a whole-brain level, we applied an *exclusive* mask constructed from the intersection of the main effect of coherence and the coherence × distance interaction (each map thresholded at *p* < .05, uncorrected) and searched for a main effect of distance (*p* < .05, FWE-corrected). This analysis identified clusters in posterior cingulate cortex, with higher activity when distance was high, and pre-supplementary motor area (pre-SMA), with higher activity when distance was low (**Fig. 3a-b**). We obtained comparable whole-brain effects when including correct trials only (**SI Appendix: Fig. S5**).

### Controlling for choice reaction time and value

We next considered alternative explanations of our neural results in terms of choice reaction time and choice value. The effects of coherence, distance and the coherence × distance interaction on pgACC activity survived the inclusion of choice reaction time, both in a regression analysis of activity time courses (**Fig. 3c**) and a series of control GLMs (**SI Appendix: Fig. S5**). We also took advantage of the fact that we varied the reward magnitude associated with the scoring rule on trials where an explicit confidence estimate was required, such that a correct decision was three times more valuable on half of these confidence trials. While reward magnitude modulated activity time courses in ventral striatum, in line with its role in encoding reward expectation (21), we did not observe an effect of reward magnitude in pgACC (**Fig. 3d**). Taken together, these analyses indicate that the neural activations identified by our factorial design were not simply due to variation in choice reaction time and/or choice value.

### Neural basis of decision confidence

Having demonstrated that pgACC tracks expected performance as specified by our factorial design, we sought to establish a role for pgACC in the construction of a subjective sense of decision confidence. Normatively, decision confidence should reflect an internal estimate of the probability that a choice is correct, and should therefore, if computed accurately, track expected performance as assayed using our factorial design. Indeed, the above analyses show that pgACC satisfies such a requirement for a neural signature of decision confidence. However, a direct correspondence between expected performance and decision confidence should not be expected, especially in the absence of any feedback: at the single-trial level, subjective confidence is an internal state that can vary even when external variables are held constant, and at the aggregate level, subjects may not have assigned appropriate weights to the components of confidence formation. To establish that pgACC is central to a subjective sense of decision confidence, it is therefore important to show that pgACC also tracks such ‘residual’ variation in subjective confidence over and above expected performance.

We first sought to establish a trial-by-trial relationship between pgACC activity and subjective confidence. We fitted an ordinal regression model to each subject’s explicit confidence estimates in the pre-scan session and used this model to generate out-of-sample predictions about their subjective confidence in the scan session (**Fig. 4a**). Supporting that pgACC is central to a subjective sense of decision confidence, pgACC activity estimates (**Fig. 4b**, linear regression, *t*(31) = 2.14, *p* = .041) and pgACC activity time courses (**Fig. 4c**) predicted trial-by-trial variation in this model-derived variable. Further, pgACC activity estimates (**Fig. 4b**, linear regression, *t*(31) = 1.26, *p* = .22) and pgACC activity time courses (**Fig. 4c**) also predicted trial-by-trial variation in the explicit confidence estimates elicited every 5-10 trials.

**Figure 4.**
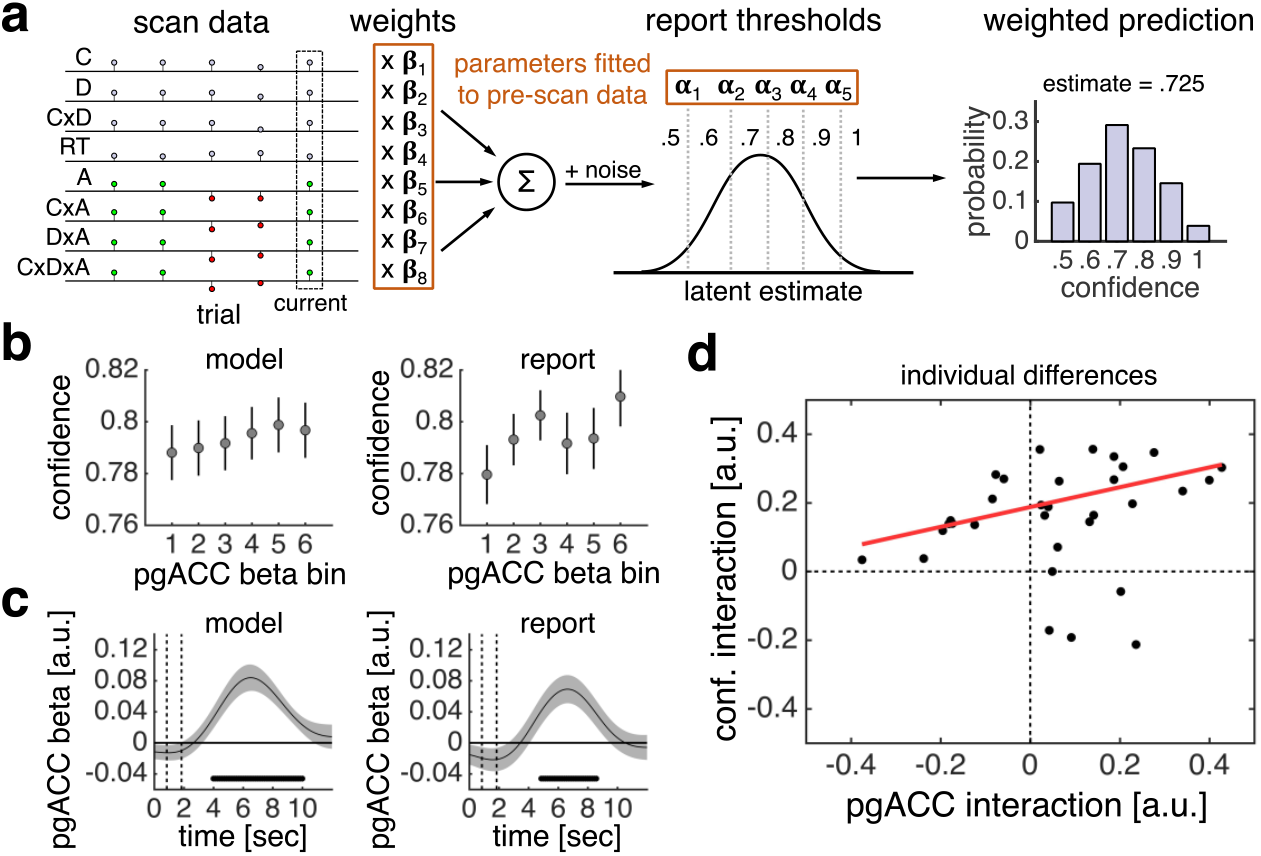
Activity in pgACC predicts decision confidence. **a**, Model of subjective confidence. We fitted an ordinal regression model to each subject’s confidence estimates in the pre-scan session. The model has a set of weights, which parameterise the effects of stimulus and choice features on confidence estimates, and a set of thresholds, which parameterise report biases. By applying the fitted model to each trial of a subject’s scan data (stimulus and choice features), we generated a prediction about their subjective confidence on that trial. The prediction is a probability distribution over possible responses (e.g., ‘.5’ has a 10% probability, ‘.6’ has a 20% probability, and so forth). We used the expectation over possible responses as our current estimate of subjective confidence. See **SI Appendix: Fig. S8** for model evaluation. C: coherence. D: distance. RT: reaction time. A: accuracy. **b**, Visualisation of encoding of model-derived subjective confidence (all trials) and reported confidence (confidence trials) in single-trial pgACC activity estimates. **c**, General linear model analysis of encoding of model-derived subjective confidence (all trials) and reported confidence (confidence trials) in pgACC activity time courses. Dots below time course indicate significant excursion of *t*-statistics assessed using two-tailed permutation tests. **d**, Correlation between interaction of coherence and distance in pgACC activity and confidence estimates. Interaction effects were calculated as a ‘difference of differences’ in our 2×2 design. The pgACC interaction was calculated using all trials; the confidence interaction was calculated using confidence trials only. We used robust linear regression as 4 outlying subjects were, on high-coherence trials, more confident when distance was low than when distance was high – a pattern which is inconsistent with normative predictions for decision confidence and the behavioural results shown in **Fig. 2c. b-c**, Data are represented as group mean ± SEM.

We next assessed the relationship between individual differences in the neural profile of pgACC activity and the behavioural profile of confidence estimates. There was substantial variation in the extent to which subjects’ explicit confidence estimates reflected an interaction between coherence and distance (**SI Appendix: Fig. S2**). Notably, the degree to which coherence and distance interacted in subjects’ pgACC activity predicted the degree to which the factors interacted in subjects’ explicit confidence estimates (**Fig. 4d**; robust linear regression, *t*(30) = 2.78, *p* = .009). This relationship survived controlling for the corresponding effect for expected performance (robust linear regression, *t*(29) = 2.44, *p* = .021), further supporting that pgACC activity is key to the construction of a subjective sense of decision confidence.

## Discussion

In studies of the neural basis of decision confidence it has proven difficult to dissociate a neural representation of decision confidence from neural representations of its component parts. For example, in the context of the classic random dot-motion task, a neural area may predict opt-out responses because it tracks the reliability of a percept of motion direction – a key component of a confidence computation – rather than decision confidence itself – a quantity which often requires the integration of multiple components (1). Recognizing this distinction is key for advancing our understanding of the neurobiology of decision confidence.

We developed a psychophysical approach to dissociate a neural representation of decision confidence from neural representations of its component parts. Extending the classic random dot-motion task, we varied both the percentage of coherently moving dots, which could move in any direction along the full circle, and the angular distance between the net direction of dot motion and a variable reference against which motion direction had to be judged. We showed that, in this design, subjects’ expected performance, an aggregate signature of decision confidence, is a function of both sensory reliability and boundary distance. Using our task to interrogate fMRI data, we observed that activity in an area in medial prefrontal cortex (pgACC) uniquely tracked expected performance, whereas neural areas previously proposed to track decision confidence tracked sensory reliability (posterior parietal cortex and ventral striatum) or boundary distance (pre-SMA). Supporting that the information encoded by pgACC is central to a subjective sense of decision confidence, we found that pgACC activity predicted both within-subject and between-subject variation in explicit confidence estimates.

Intriguingly, pgACC may be involved in the formation of not only a local estimate of probability correct on a single trial, the computational definition of decision confidence, but also a more global estimate of probability correct across trials, an estimate which is critical for assessing one’s general ability on a task. Evidence for the latter function comes from a recent study which showed that pgACC activity tracks a running average of subjects’ performance history and predicts their explicit evaluations of expected performance, independently of the expected value of a trial, as in our study (22). Unlike our study in which performance was designed to be independent from one trial to another, performance was rigged such that it was autocorrelated across trials and could only be learnt by tracking performance feedback. The diverse connectivity profile of pgACC is consistent with this area as central to the formation of both local and global estimates of probability correct across different task domains: pgACC is connected to surrounding medial prefrontal cortex, dorsolateral and ventrolateral prefrontal cortex, cingulate cortex, subcortical areas, such as hippocampus and striatum, and posterior areas, including parietal cortex (23). Taken together, these findings may help explain why metacognition, the ability to monitor and evaluate the success of one’s task performance, is often impaired after prefrontal lesions and/or degeneration (24). If pgACC is critical for confidence formation, compromising pgACC, or connections to and/or from pgACC, should naturally lead to discrepancies between subjective evaluations and objective performance.

It is noteworthy that our factorial analysis did not identify rostrolateral prefrontal cortex (rlPFC). Previous studies have consistently shown that rlPFC activity tracks explicit confidence estimates (11, 12), and that the microstructure of rlPFC predicts the degree to which an individual’s subjective evaluations reflect objective performance (25). One hypothesis, which may reconcile these results with ours, is that rlPFC is not itself involved in computing an internal estimate of the probability that a choice is correct, but instead governs the mapping of this variable onto an explicit confidence estimate for report. In support of this hypothesis, we found that activity time courses in an rlPFC ROI also predicted trial-by-trial variation in the model-derived predictor of subjective confidence and subject’s explicit confidence estimates (**SI Appendix: Fig. S9**). There is evidence that rlPFC manages task sets and rules (26), functions which presumably are involved in maintaining a consistent confidence mapping, or updating a confidence mapping in response to a particular communicative context (4). Future studies directly manipulating confidence mappings are required to test this hypothesis.

Several studies have proposed that decision confidence is a function of the absolute distance between sensory evidence, *x*, and a choice boundary, *b*: |*x* – *b*| (3, 6, 8, 9). This formulation makes the prediction that decision confidence is, on correct trials, higher for larger distances but, on error trials, lower for larger distances – a qualitative pattern which is mirrored in the activity of putative neural substrates of decision confidence, such as rat orbitofrontal cortex (3). Interestingly, we found that pre-SMA, which tracked boundary distance in our task, also showed such an activity pattern (**SI Appendix: Fig. S10**). This area has been implicated in conflict monitoring (27), confidence formation (12) and changes of mind (28). A unifying account of these results may be that pre-SMA encodes evidence in the coordinate frame of response options – a signal which can be used to guide subsequent behaviour and cognition, such as increasing response caution (29) and increased weight to post-choice evidence (28). However, in our task, decision confidence is a function of not only boundary distance but also sensory reliability – as expected under normative models of decision confidence and as shown by subjects’ explicit confidence estimates (**Fig. 2c**).

Our study may prompt a reconsideration of the contribution of intraparietal sulcus (IPS) and ventral striatum to confidence-based behaviours. For example, in the context of the classic random dot-motion task, neurons in IPS, including lateral intraparietal area (LIP) which receives inputs from motion-sensitive area MT+ (30), have been shown to encode the accumulation of evidence towards a choice boundary, with the amplitude and the temporal profile of neuronal activity varying with motion coherence and choice reaction time (31). Because such activity patterns carry information about the probability that a choice is correct, IPS has been proposed to be central to not only choice but also confidence formation (5). A similar argument has been made for dopaminergic neurons in substantia nigra which are connected with ventral striatum (8). However, in the context of our continuous-direction, variable-reference random dot-motion task, we observed that fMRI activity in a putative human homologue of IPS and ventral striatum tracked sensory reliability and not the integration of sensory reliability with boundary distance. These results are in line with a recent study which found that superior colliculus, which together with IPS is also part of the oculomotor planning circuit, tracks choice and not confidence formation (32).

We remain agnostic as to the source of these reliability-related signals in our task. However, our results are consistent with a hypothesis that such areas encode a probability distribution over sensory states given sensory evidence (33). On this account, the amplitude of neural activity in these areas may reflect the reliability associated with this distribution. First, activity in IPS and ventral striatum cannot be explained by a bottom-up response to coherent motion but instead appears to track *changes* in motion coherence: in a separate motion-localiser scan we found that extrastriate cortex, including MT+, but notably not IPS or ventral striatum, was activated in a contrast between coherently moving and static dots (**SI Appendix: Fig. S11**). Second, activity in these areas is *selective* for changes in motion coherence as evidenced by our masking approach (**Fig. 3a**). Finally, subjects’ percepts of motion direction were more reliable on high-coherence trials (**Fig. 2a**), and estimates of sensory reliability were clearly used to inform confidence estimates (**Fig. 2c**). An alternative interpretation is that activity in IPS and/or ventral striatum reflects expected reward across possible choices (33), a quantity which would be larger on high-coherence trials. It remains to be seen how activity in these areas varies with the space of sensory states (e.g., binary versus continuous direction) and the onset of the choice boundary (e.g., concurrent versus delayed).

Why should the brain maintain both ‘choice-dependent’ and ‘choice-independent’ estimates of certainty? In many tasks, to perform optimal inference, it is useful to represent the certainty associated with relevant sensory or cognitive variables independently of any future choice (1, 34). For example, when inferring the length of an object from visual and haptic information, the brain needs to know which source is more reliable and therefore should dominate the integrated visual-haptic percept, regardless of which response may eventually be required (35). However, after making a choice, it may be efficient to combine these estimates into a statistic summarising the probability that a choice is correct (1), which can be used to guide multi-stage decisions, control learning from feedback (36) and optimise group decisions (4). Our study supports a distinction between choice-dependent and choice-independent estimates of certainty (1), and indicates a dissociation in the neural encoding of these quantities.

## Materials and Methods

### Subject details

Thirty-five adults (17 female, mean ± SD age = 23.60 ± 4.31 years) with normal or corrected-to-normal vision participated in the study. Subjects performed separate pre-scan and scan sessions (2-14 days between sessions). Three subjects were excluded due to excessive motion and/or sleeping during the scan session; their data was not included in any analyses. Subjects provided informed consent, and the study was approved by the ethics committee of University College London. Subjects received a flat rate for participation (£40) and earnt an additional performance-based bonus (up to £12).

### Experimental details

#### Task

Subjects performed a continuous-direction, variable-reference random dot-motion task as described in the text (**Fig. 1**). Choices and confidence estimates were submitted to a variant of the Brier score, with a subject’s score (reward or loss) on trial *t* calculated as: *score*_t_ = *r* × (.5 – *(confidence*_t_ – *accuracy*_t_)^2^), where *confidence*: {.5, .6, .7, .8, .9, 1}, *accuracy:* {0, 1}, and *r* is a scaling factor which specifies the maximum reward or cost. In the pre-scan session, the reward factor was fixed at £4. In the scan session, the reward factor was £2 on half of confidence trials and £6 on the other half (indicated by ‘£’ or ‘££’ displayed above the scale). Subjects received the sum of their average trial-by-trial earnings calculated separately for each reward factor (maximum: £12). See **SI Appendix: Methods** for details about trial events and timings, response mode and stimulus presentation.

#### Pre-scan procedure

The pre-scan session consisted of five runs. Subjects first viewed motion stimuli of variable durations and variable coherences. Subjects then practiced the task (40 trials) with high coherence and high distances. In this run only, subjects received trial-by-trial feedback, with the aim to familiarise subjects with making direction judgements in continuous space. Subjects then performed calibration phases 1 (120 trials) and 2 (260 trials) in which we estimated a set of coherences and distances (2×4 design) associated with target levels of choice accuracy. Finally, subjects performed the main experiment (540 trials). See **SI Appendix: Methods** for details about calibration procedures.

#### Scan procedure

The scan session comprised seven runs. Subjects first performed a calibration phase during the acquisition of structural images (180 trials). Subjects then performed the main experiment over five runs (5×112 = 560 trials). We used a subset of the calibrated task parameters (2×2 design) from the pre-scan session. In the final scan run, subjects viewed alternating displays (12s) of static and dynamic dots (2×12 = 24 displays). See SI Appendix: Methods for details about calibration procedure.

### Behavioural analysis

#### Choice reaction time

We excluded trials on which subjects’ choice reaction times were 2.5 SD below or above their grand mean reaction time computed separately for the pre-scan and the scan sessions. This procedure resulted in the exclusion of approximately 2% of trials per subject per session.

#### Regression models

We used logistic regression to predict choice accuracy, linear regression to predict choice reaction time and ordinal regression to predict confidence estimates (**Fig. 2**). We log-transformed choice reaction time, contrast-coded coherence and distance, and z-transformed all variables. We performed a separate regression for each subject and tested the group-level significance of a predictor by comparing the coefficients pooled across subjects to zero (one-sample *t*-test). We used ordinal regression to construct the computational model of subjective confidence for fMRI analysis (**Fig. 4a**). The model included eight predictors: log-transformed choice reaction time, accuracy, coherence, distance, and interaction terms for coherence × distance, accuracy × coherence, accuracy × distance and accuracy × coherence × distance. We did not z-score variables to facilitate between-session compatibility. The model provided a good fit to the pre-scan data and generalised well to the scan data (**SI Appendix: Fig. S8**).

#### Hierarchical drift-diffusion model

We employed hierarchical Bayesian estimation of subjects’ drift-diffusion model (DDM) parameters using the HDDM toolbox (17) (http://ski.clps.brown.edu/hddm_docs/). We fitted drift-rate (v), decision threshold (*a*) and non-decision time (*t*) separately for each condition of our factorial design (pre-scan: 8 conditions; scan: 4 conditions). We also included inter-trial variability in drift-rate (sv) and non-decision time (st) as free parameters that were constant across conditions. We extracted mean group-level posterior estimates for visualisation of DDM parameters (**SI Appendix: Fig. S4**) and entered these into the *simuldiff* function from the DMAT toolbox (37) (http://ppw.kuleuven.be/okp/software/dmat/) to generate posterior predictives (**Fig. 2a-b**).

### FMRI analysis

#### Whole brain

FMRI analysis was conducted using SPM 12 (www.fil.ion.ucl.ac.uk/spm/). The whole-brain analysis shown in **Fig. 3a** was based on a single event-related General Linear Model (GLM1). We labelled each trial according to whether Coherence (C) and Distance (D) was Low (L) or High (H): CL&DL, CL&DH, CH&DL, CH&DH. We modelled the four trial types with separate ‘condition’ regressors. Choice reaction-time outlier trials were assigned to an ‘outlier’ regressor. These regressors were specified as boxcars time-locked to the onset of the motion stimulus and spanning until choice (38). We modelled confidence events with a separate ‘rate’ regressor, specified as a stick function time-locked to the onset of the scale and parametrically modulated by the (z-transformed) number of button presses used to navigate the marker. We included motion and biophysical parameters as additional ‘nuisance’ regressors. Regressors were convolved with a canonical hemodynamic response function. Regressors were modelled separately for each scan run and constants were included to account for between-run differences in mean activation and scanner drifts. A high-pass filter (128s cut-off) was applied to remove low-frequency drifts.

We assessed group-level significance by applying one-sample *t*-tests against zero to the first-level contrast images. Inclusive/exclusive masks for our factorial analysis were created using SPM’s *imcalc* function; each second-level contrast image was thresholded at *p* < .05, uncorrected, the default setting for masking analyses in SPM, before creating a mask. We report clusters significant at *p* < .05, FWE-corrected for multiple comparisons, with a cluster-defining threshold of *p* < .001, uncorrected.

We obtained similar results in GLMs that introduced modifications to GLM1 (**Fig. S5**). In GLM1C, only correct trials were included. In GLM2, the four condition regressors had a fixed duration of 1s and were parametrically modulated by (log-transformed) choice reaction time. In GLM2C, only correct trials were included. In GLM3, there was only one ‘interest’ regressor which had a fixed duration of 1s and which was parametrically modulated by four dummy variables, one for each trial type, and (log-transformed) choice reaction time. In GLM3C, only correct trials were included. Parametric modulators were not orthogonalized and regressors competed to explain variance.

Clusters surviving whole-brain correction post- and pre-masking for GLM1 and GLM1C are shown in **SI Appendix: Tables S1-S4**. See **SI Appendix: Methods** for details about fMRI acquisition, preprocessing and physiological monitoring.

#### Regions of interest

See **SI Appendix: Methods** for details about ROI specification, ROI single-trial activity estimates, ROI activity time courses and permutation testing.

### Resource sharing

Anonymised behavioural data and code supporting main analyses are available on GitHub (https://github.com/metacoglab/BangFleming/). Unthresholded group-level statistical maps are available on NeuroVault (https://neurovault.org/collections/3792/).

## Acknowledgements

We thank Chris Summerfield for helpful comments on an earlier version of this manuscript, and Dick Passingham for advice on neuroanatomy. The Wellcome Centre for Human Neuroimaging is supported by core funding from the Wellcome Trust (203147/Z/16/Z).

## Supporting Information: Appendix

### Methods

#### Task

##### Trial events and timings

Each trial began with the presentation of a fixation cross at the centre of a circular aperture. After a uniformly sampled delay (pre-scan: .5-1s; scan: 1-4s), subjects viewed a field of moving dots (1s). Subjects were instructed to fixate during stimulus presentation. Once the motion stimulus terminated, subjects were presented with a direction reference which transected the aperture. Subjects were required to press one of two buttons to indicate whether the net direction of dot motion (angle as measured from aperture centre) was clockwise (CW) or counter-clockwise (CCW) to the reference (angle as measured from aperture centre). As a visual aid, the response buttons and the associated arcs of the aperture were coloured orange and blue, with colour assignment counterbalanced across subjects. Once a choice had been made, the colour of the central cross (orange or blue) confirmed the decision (.25s). In the pre-scan session, after having made a choice, subjects were asked to estimate the confidence in their choice on every trial. In the scan session, subjects were asked to estimate the confidence in their choice every 5-10 trials (15% of trials). Subjects responded by moving a marker along a scale from 50% to 100% in steps of 10%. The marker started randomly in one of the six locations along the scale and was controlled by button press. Once a response had been registered, the marker turned grey (.5s), before the next trial started.

##### Motion stimulus

The motion stimulus was made up of three sets of dots (each dot was 0.12 degrees in diameter) shown in consecutive frames inside the circular aperture (8 degrees in diameter) centred on the fixation cross (0.2 degrees in diameter). Each set of dots was shown for one frame (about 16ms) and then replotted again three frames later (about 50ms) – some dots were displaced in the specified motion direction at a speed of 2 degrees s^-1^ while the rest of the dots were displaced at random locations within the aperture. We refer to the percentage of dots displayed in the specified motion direction as coherence, *k*. The dot density was fixed at 16 dots degrees^-2^ s^-1^. To help subjects maintain fixation, a circular region (0.7 degrees in diameter) at the centre of the aperture was kept free of dots. The motion direction was sampled uniformly from the range 1-360 degrees. The direction of the reference (0.8 degrees in length and 0.08 degrees in width) was within 45 degrees of the motion direction. We refer to the difference between the motion direction and the direction reference as signed distance, *δ,* with positive values for CW and negative values for CCW. A set of coherences, **K**, and a set of signed distances, **Δ**, were calibrated for each subject and crossed in a factorial design in each session.

##### Pre-scan calibration

The aim of the calibration procedure was to identify a pair of coherences associated with different levels of sensory reliability and a set of signed distances associated with different levels of boundary difficulty. This aim was achieved in two phases. We did not elicit confidence estimates in either phase. In phase 1, subjects performed the task with a fixed absolute distance, **Δ**: {-10,10}. Coherence was .20 for the first two trials and then updated using a 2-down-1-up procedure: after two correct decisions, coherence was decreased by .01; after one incorrect decision, coherence was increased by .01. We used the median coherence of the last 20 trials of phase 1 to specify a medium coherence, *k*_med_, constrained to a lower limit of .12 and an upper limit of .50. In phase 2, subjects performed the task with coherence fixed at *k*_med_. We employed a range of signed distances, **Δ**: {-45, −24, −12, −6, −3, 3, 6, 12, 24, 45}, and fitted a psychometric function for each subject using logit regression. This procedure provided us with a measure of a subject’s choice bias and sensitivity. We corrected the psychometric function for choice bias and used the corrected psychometric function to infer a set of positive distances associated with the target accuracies 60%, 72.5% and 85%: *δ*_60_, *δ*_72.5_, *δ*_85_. For the main experiment, we defined the set of coherences as, **K**: {*k*_low_ = .5 × *k*_med_, *k*_high_ = 2 × *k*_med_}, and the set of signed distances as, **Δ**: {-45, -*δ*_85_, -*δ*_72.5_, -*δ*_60_, *δ*_60_, *δ*_72.5_, *δ*_85_, 45}. The extreme signed distances were probed three times less often; they served as anchor points for construction of psychometric functions. Pre-scan calibration data are shown in **Fig. S1**.

##### Scan calibration

Subjects performed one further calibration phase at the start of the scan session during the acquisition of structural images. Coherence was fixed at *k*_med_ and we calibrated the signed distances associated with 60% and 85% accuracy, ±*δ*_60_ and ±*δ*_85_, using a QUEST procedure (1). For the main experiment, we defined the set of coherences as **K**: {*k*_low_, *k*_high_} and the set of signed distances as **Δ**: {-*δ*_85_, -*δ*_60_, -*δ*_60_, *δ*_85_}. Scan calibration data are shown in **Fig. S1**.

##### Pre-scan procedure

The pre-scan session consisted of five runs. Subjects first completed a tutorial, with motion stimuli shown for variable durations, 1-5s, and variable coherences, **K**: {.30, 60}. Subjects then practiced the task (40 trials) at high coherence, **K**: {.30, .60}, and high absolute distance, **Δ**: {-30, 30}. In this run only, subjects received trial-by-trial feedback (central cross briefly turned green after correct choices and red after incorrect choices), with the aim to familiarise subjects with direction judgements in continuous space. Next, subjects performed calibration phases 1 (120 trials) and 2 (260 trials). Finally, subjects performed the main experiment (540 trials). The pre-scan session lasted 2 hours.

##### Scan procedure

The scan session was made up of seven runs. Subjects first performed the calibration phase during the acquisition of structural images (180 trials) and then the main experiment over five runs (5 × 112 = 560 trials). In the scan session only, sampling of motion direction was yoked such that there was a 50% probability that a given trial had the same motion direction as on the previous trial. This manipulation is orthogonal to coherence and distance, and direction does not tell subjects whether the upcoming choice should be CW or CCW. Effects of repetition on direction-specific neural responses will be analysed in a separate paper. In the final scan run, subjects viewed alternating displays (12s) of static and dynamic dots, *k* = .50 (2 × 12 = 24 displays).

#### Hierarchical drift-diffusion model

Decision formation was modelled using the drift-diffusion model (DDM). The DDM models two-choice decision-making as a process of accumulating noisy evidence over time with a certain speed, or drift-rate (v), until one of two decision thresholds is crossed and the associated response is executed. Larger threshold separation (a) leads to slower responses but more accurate responding. The DDM includes a non-decisional component (t) which captures time needed for stimulus encoding and motor execution. We employed hierarchical Bayesian estimation of subjects’ DDM parameters using the HDDM toolbox (2) (http://ski.clps.brown.edu/hddm_docs/). We fitted non-decision time (t), decision threshold (t) and drift-rate (v) separately for each condition of our factorial design (pre-scan: 8 conditions; scan: 4 conditions). We also included inter-trial variability in drift-rate (sv) and non-decision time (st) as free parameters that were constant across conditions. The HDDM toolbox applies Markov Chain Monte-Carlo (MCMC) sampling methods to approximate posterior distributions over the DDM parameters; we ran 1 chain with 5000 samples, with the first 1000 samples discarded as burn-in. We extracted mean group-level posterior estimates for visualisation of DDM parameters in each condition (**Fig. S4**) and generating posterior predictives (**Fig. 2a-b**). We used the *simuldiff* function from the DMAT toolbox (3) (http://ppw.kuleuven.be/okp/software/dmat/) to generate posterior predictives: for each condition, we simulated 100,000 trials under the hierarchically estimated DDM parameters and calculated mean choice accuracy and mean choice reaction time in that condition.

#### FMRI

##### Acquisition

MRI data were acquired on a 3T Siemens Trio scanner with a 32-channel head coil. T1-weighted structural images were acquired using a 3D MDEFT sequence: 1×1×1 mm resolution voxels; 176 sagittal slices, 256×224 matrix; TR = 10.55ms; TE = 3.14ms; TI = 680ms. BOLD T2*-weighted functional images were acquired using a gradient-echo EPI pulse sequence: 3×3×3 mm resolution voxels; 48 transverse slices, 64×74 matrix; TR = 3.36; TE = 30ms; slice tilt = 0 degrees, slice thickness = 2 mm; inter-slice gap = 1mm; ascending slice order. Field maps were acquired using a double-echo FLASH (gradient echo) sequence: TE1 = 10ms; TE2 = 12.46ms; 64 slices were acquired with 2 mm slice thickness and a 1 mm gap; in-plane field of view is 192×192 mm^2^ with 3×3 mm^2^ resolution.

##### Pre-processing

MRI data were pre-processed using SPM12 (Wellcome Trust, London). The first 4 volumes of each functional run were discarded to allow for T1 equilibration. Functional images were slice-time corrected, realigned and unwarped using the field maps (4). Structural T1-weighted images were co-registered to the mean functional image of each subject using the iterative mutual-information algorithm. Each subject’s structural image was segmented into grey matter, white matter and cerebral spinal fluid using a nonlinear deformation field to map it onto a template tissue probability map (5). These deformations were applied to structural and functional images to create new images spatially normalised to the Montreal Neurological Institute (MNI) space and interpolated to 2×2×2 mm voxels. Normalized images were spatially smoothed using a Gaussian kernel with full-width half-maximum of 8mm. The motion correction parameters estimated from the realignment procedure and their first temporal derivatives – 12 ‘motion’ regressors in total – were included as confounds in the first-level analysis for each subject.

##### Physiological monitoring

Peripheral measurements of a subject’s pulse and breathing were made together with scanner slice synchronisation pulses using a Spike2 data acquisition system (Cambridge Electronic Design Limited, Cambridge UK). The cardiac pulse signal was measured using an MRI compatible pulse oximeter (Model 8600 F0, Nonin Medical, Inc. Plymouth, MN) attached to a subject’s finger. The respiratory signal, thoracic movement, was monitored using a pneumatic belt positioned around the abdomen close to the diaphragm. A physiological noise model was constructed to account for artifacts related to cardiac and respiratory phase and changes in respiratory volume using an in-house MATLAB toolbox (6). Models for cardiac and respiratory phase and their aliased harmonics were based on RETROICOR (7) and a similar, earlier method (8). Basis sets of sine and cosine Fourier series components extending to the 3rd harmonic were used to model physiological fluctuations. Additional terms were included to model changes in respiratory volume (9, 10) and heart rate (11). This procedure yielded a total of 14 ‘biophysical’ regressors which were sampled at a reference slice in each image volume. The regressors were included as confounds in the first-level analysis for each subject.

##### Regions of interest

ROI masks for bilateral IPS, pre-SMA and pgACC were created by applying a leave-one-out procedure to activations from GLM1. In particular, for each subject, we performed second-level analyses excluding the subject’s data and used the resulting second-level t-maps (cluster-defining threshold: p < .001, uncorrected) to specify the subject’s ROI masks. This ROI procedure ensured unbiased ROI selection as each subject’s neural activity did not contribute to the selection of ROIs for that subject. We used clusters associated with a main effect of coherence (bilateral IPS), a main effect of distance (pre-SMA), and the coherence × distance interaction (pgACC). ROI masks for MT+ were based on the localiser scan; we created a bilateral group mask using the second-level contrast between dynamic and static motion, and then, for each subject, created bilateral MT+ masks (8-mm sphere) around the subject-specific peaks inside the bilateral group mask. ROI masks for bilateral ventral striatum were specified using the Oxford-GSK-Imanova Striatal Structural atlas included with FSL. ROI masks for bilateral rostrolateral prefrontal cortex were specified using the maps provided by Neubert and colleagues (12) (union of ‘46’ and ‘fpl’).

##### Activity time courses

We transformed each ROI mask from MNI to native space and extracted pre-processed BOLD time courses as the average of voxels within the mask. For each scan run, we regressed out variation due to head motion and biophysical responses, applied a high-pass filter (128s cut-off) to remove low-frequency drifts, and oversampled the BOLD time course by a factor of ~23 (time resolution of .144s). For each trial, we extracted activity estimates in a 12s window (84 time points), time-locked to 1s before the onset of the motion stimulus or 2s before the onset of the confidence scale (only for reward-magnitude analysis). We used linear regression to predict ROI activity time courses. More specifically, we applied a linear regression to each time point and then, by concatenating beta-weights across time points, created a beta-weight time course for each predictor of a regression model. We performed this step separately for each subject and pooled beta-weight time courses across subjects for visualisation. We tested group-level significance using a permutation procedure. We repeated the above steps 1,001 times, for each repetition shuffling a subject’s trial time courses, and then asked, at each time point, whether the *t*-statistics associated with the empirically observed group-level effect (one-sample *t*-test of beta-weights pooled across subjects against zero) was smaller or larger than 97.5% of the *t*-statistics obtained from the permutation procedure.

##### Single-trial activity estimates

We estimated single-trial ROI activity estimates as a beta timeseries. This was achieved by means of an event-related model with a separate regressor for each trial. Regressors were boxcars time-locked to the onset of the motion display and spanning 1 second (**Fig. 4b**) or until choice (**Fig. 4d**) (13). Each regressor was convolved with a canonical hemodynamic response function. We included motion and biophysical parameters as ‘nuisance’ regressors. Regressors were modelled separately for each scan run and constants were included to account for between-run differences in mean activation and scanner drifts. A high-pass filter (128s cut-off) was applied to remove low-frequency drifts. One important consideration in using single-trial activity estimates is that a beta for a given trial can be strongly affected by acquisition artefacts that co-occur with that trial (e.g., scanner pulse artefacts). Therefore, for each subject, we computed the grand-mean beta estimate across both voxels and trials and excluded any trial whose mean beta estimate across voxels was 3 SDs below or above this grand mean (14). About 2% of trials were excluded per subject. Finally, we used the ROI masks to extract ROI single-trial activity estimates.

---

**Figure S1.**
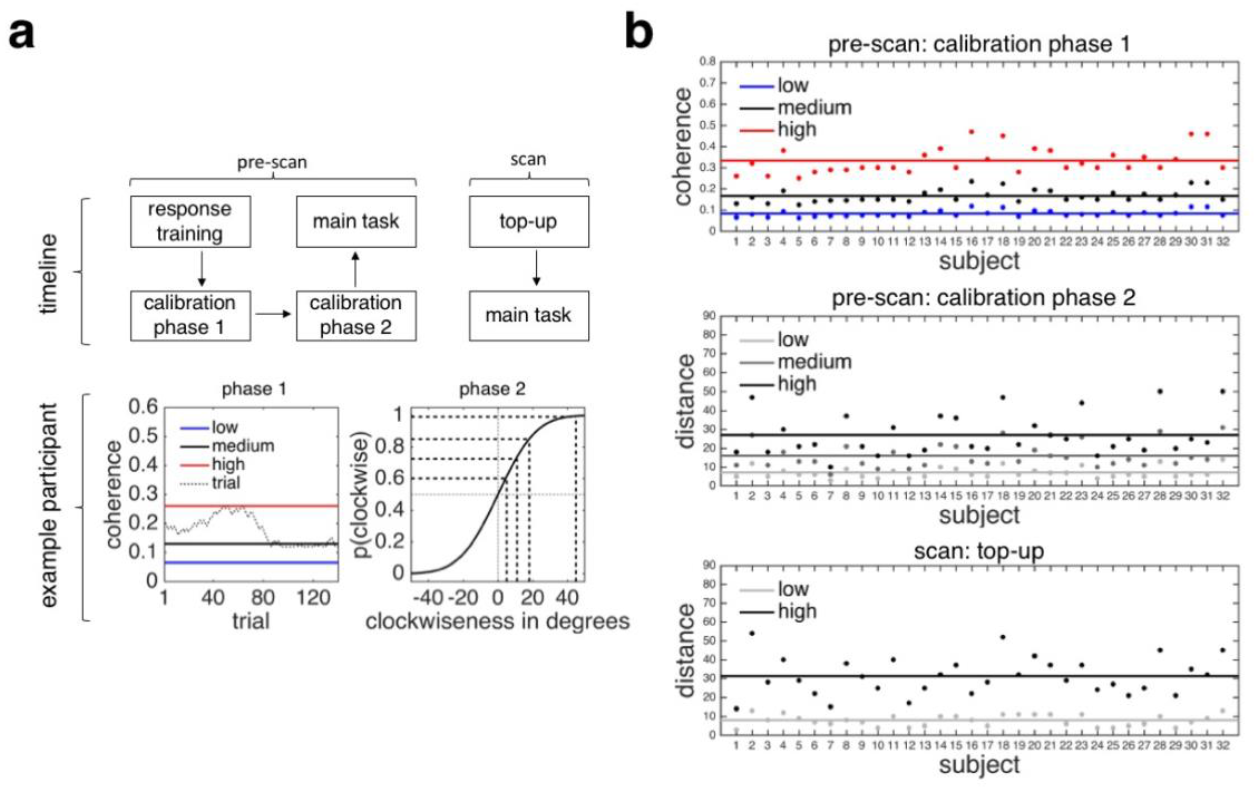
Stimulus calibration, **a**, Experimental procedure. *(top)* Subjects took part in separate pre-scan and scan sessions (2-14 days between sessions). In pre-scan calibration phase 1, we calibrated coherence (medium) at a fixed distance for each subject using a 2-down-1-up procedure. We used 50% (low) and 200% (high) of the medium coherence value in the main task, both in the pre-scan and in the scan session. In pre-scan calibration phase 2, we presented each subject with a range of distances at medium coherence so as to construct their psychometric function. We extracted a set of distances associated with target levels of accuracy for the pre-scan main task. In the scan top-up session, using an adaptive QUEST procedure, we re-calibrated a subset of distances from the pre-scan session at medium coherence for the scan main task. *(bottom)* Example data from pre-scan calibration phase 1 and 2. **b**, Calibration results. Calibrated stimulus values are shown for *(top)* pre-scan calibration phase 1, *(middle)* pre-scan calibration phase 2 and *(bottom)* scan top-up.

**Figure S2.**
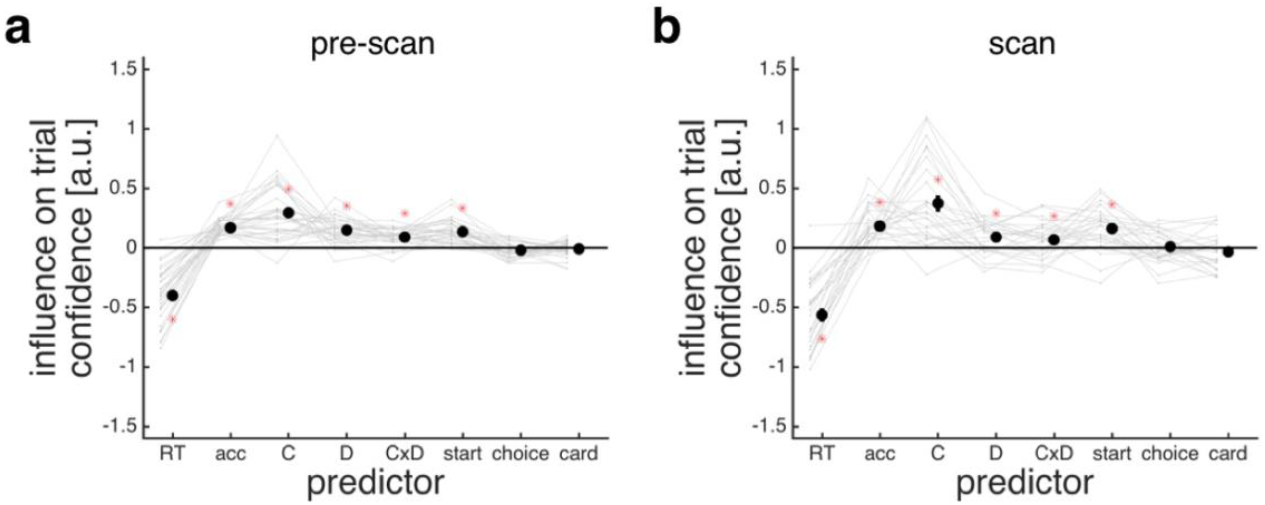
Extended regression analysis of predictors of confidence estimates. **a**, Pre-scan session. Coefficients (y-axis) from a trial-by-trial ordinal regression testing the influence of stimulus and choice features (x-axis) on current confidence. Predictors included: choice reaction time (RT), choice accuracy (acc), coherence (C), distance (D), the interaction between coherence and distance (CxD), the initial position of the confidence marker (start), whether the choice was clockwise (choice), and the cardinality of the motion direction (card). **b**, Scan session. Same regression analysis as in panel A, except that we also included the reward factor and whether the motion direction was the same as on the previous trial; these predictors were not significant and are not shown here. Confidence estimates were elicited every 5-10 trials (84 trials in total) in the scan session. **a-b**, We performed a regression for each subject and tested significance (red asterisk) by comparing the coefficients pooled across subjects to zero (one-sample *t*-test). We log-transformed choice reaction time and z-transformed predictors. Faint lines connect the coefficients for each subject. Data are represented as group mean ± SEM.

**Figure S3.**
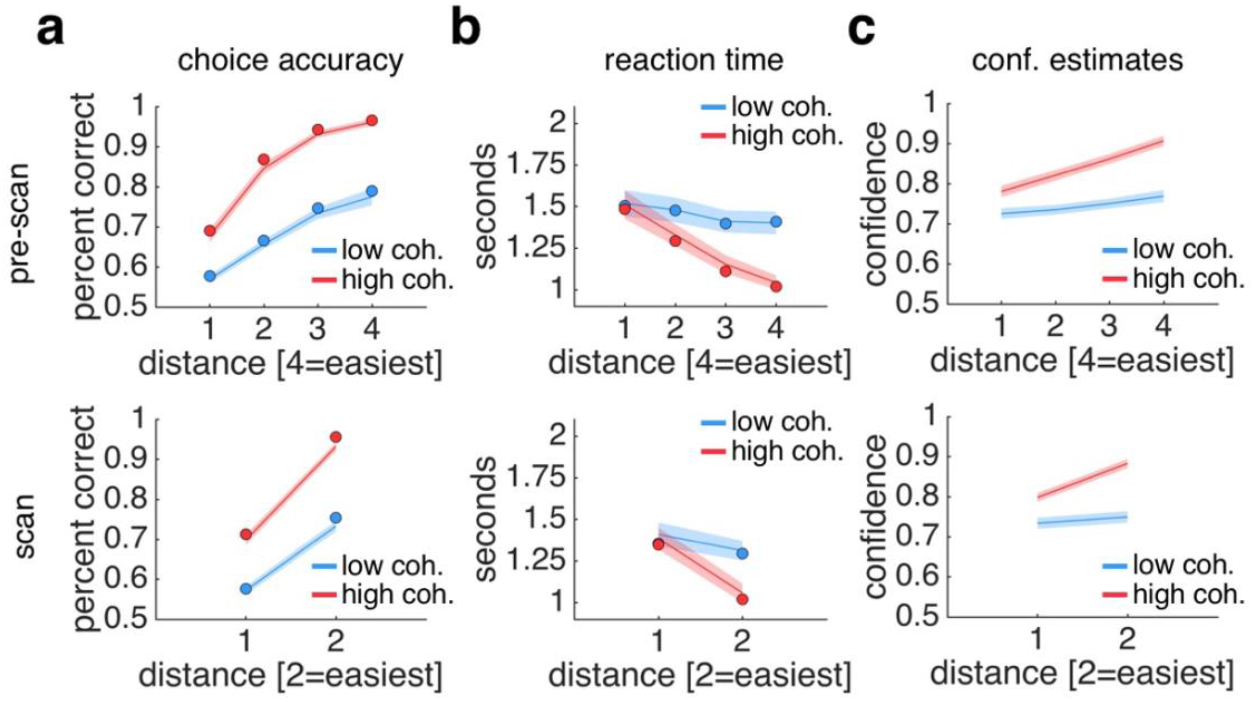
Behavioural results and HDDM fits from pre-scan and scan sessions. **a**, Choice accuracy. In the scan session, subjects were more likely to be correct when coherence was high and when distance was high (logistic regression, coherence: *t*(30) = 13.61, *p* < .001, distance: t(30) = 14.76, *p* < .001, interaction: t(30) = 11.48, *p* < .001). **b**, Reaction time measured from reference onset. In the scan session, subjects made faster decisions when coherence was high and when distance was high (linear regression, coherence: t(30) = −6.37, *p* < .001, distance: t(30) = −9.66, *p* < .001, interaction: t(30) = – 8.65, *p* < .001). **c**, Confidence estimates. In the scan session, subjects were more confident when coherence was high and when distance was high (ordinal probit regression, coherence: t(30) = 6.22, *p* < .001, distance: t(30) = 5.87, *p* < .001, interaction: t(30) = 4.63, *p* < .001). **a-b**, Solid dots are posterior predictives from the hierarchical drift-diffusion model fit to subjects’ responses separately for each condition. **a-c**, See main text for test statistics for pre-scan session. Data are represented as mean ± SEM.

**Figure S4.**
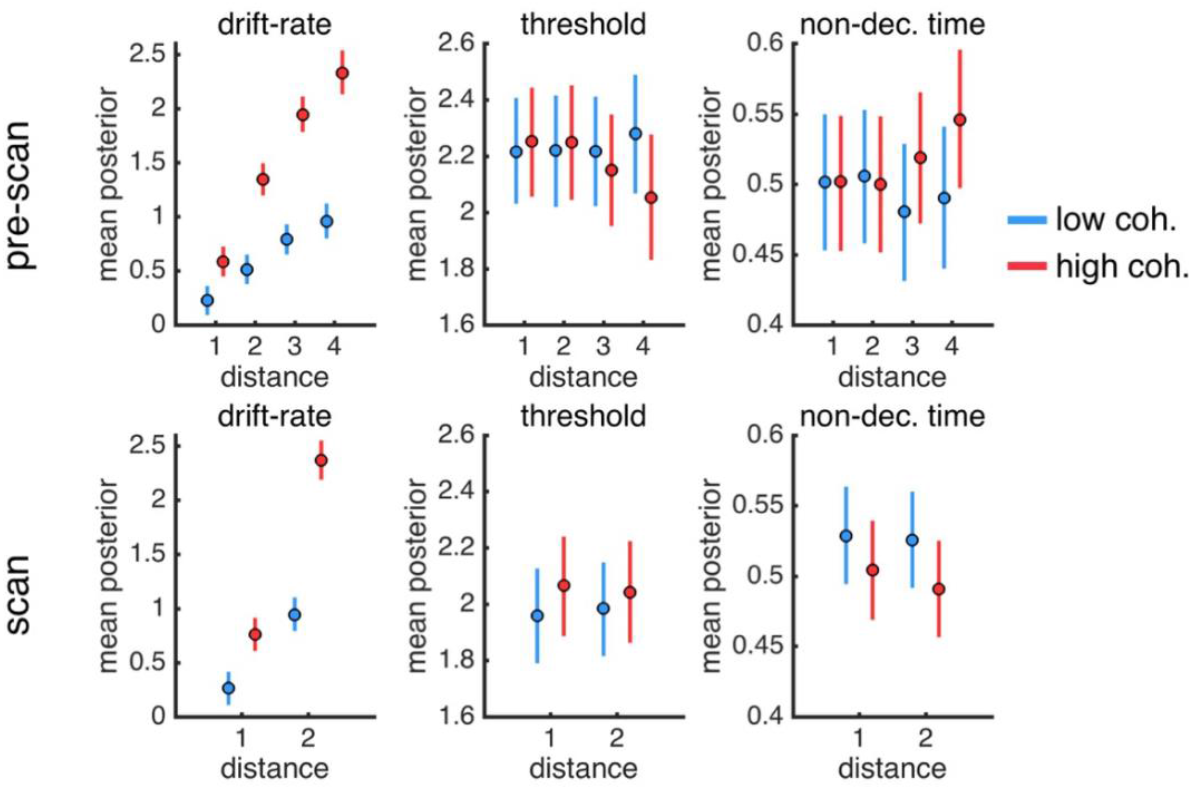
Drift-diffusion model. Mean posterior estimates obtained from hierarchical Bayesian estimation of subjects’ drift-diffusion model parameters using the HDDM toolbox for *(top)* the pre-scan session and *(bottom)* the scan session. We fitted drift-rate (v), decision threshold (a) and non-decision time (*t*) separately for each condition of our factorial design (pre-scan: 8 conditions; scan: 4 conditions). We also included inter-trial variability in drift-rate (*sv*) and non-decision time (*st*) but these parameters were fitted across conditions (pre-scan, *sv* = 0.904, *st* = 0.232; scan, *sv* = 1.063, *st* = 0.270). Error bars indicate 95%-confidence intervals as estimated from the posterior distributions over parameters.

**Figure S5.**
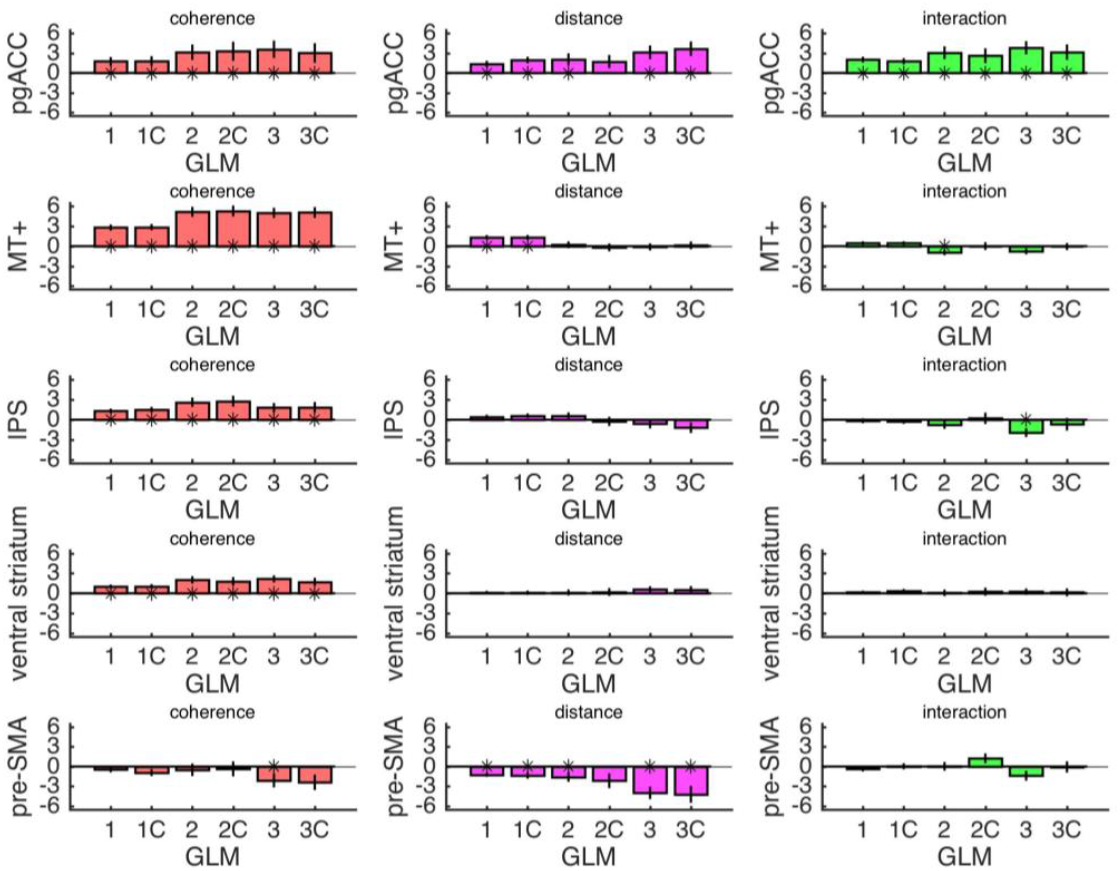
ROI contrast estimates from whole-brain control models. The plots show contrast estimates (y-axis) for coherence, distance and the coherence × distance interaction extracted from a set of control models (x-axis). We tested significance by comparing the contrast estimates pooled across subjects against zero (one-sample *t*-test). The rationale behind the three families of GLMs is as follows: GLM1s assume that choice reaction time is not a confound but reflects relevant neural processing; GLM2s control for within-condition variation in choice reaction time; and GLM3s control for between-condition variation in choice reaction time. C: correct trials only. Data are represented as group mean ± SEM.

**Figure S6.**
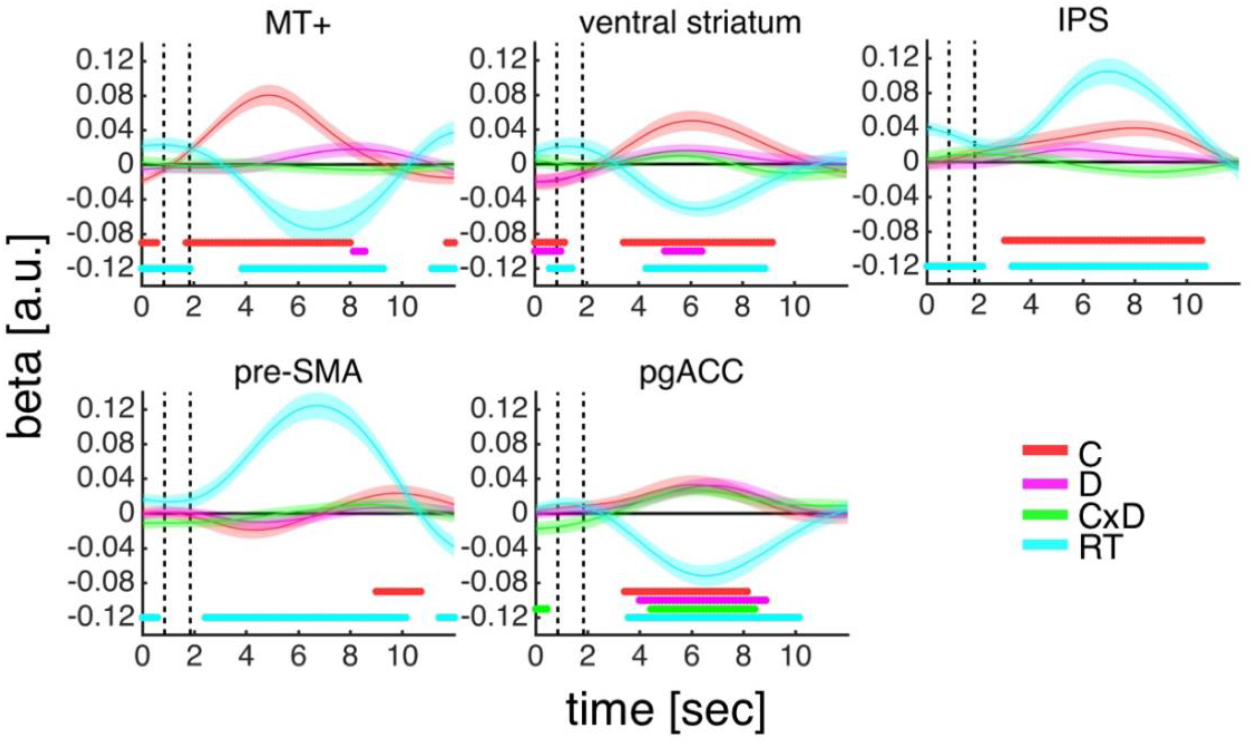
Encoding of factors in ROI activity time courses. General linear model analysis of the effects of coherence (C), distance (D), the coherence × distance interaction (CxD) and choice reaction time (RT) on ROI activity time courses. Dots below time course indicate significant excursion of *t*-statistics assessed using two-tailed permutation tests. Vertical dashed lines indicate the onsets of the motion stimulus and the choice phase. Data are represented as group mean ± SEM.

**Figure S7.**
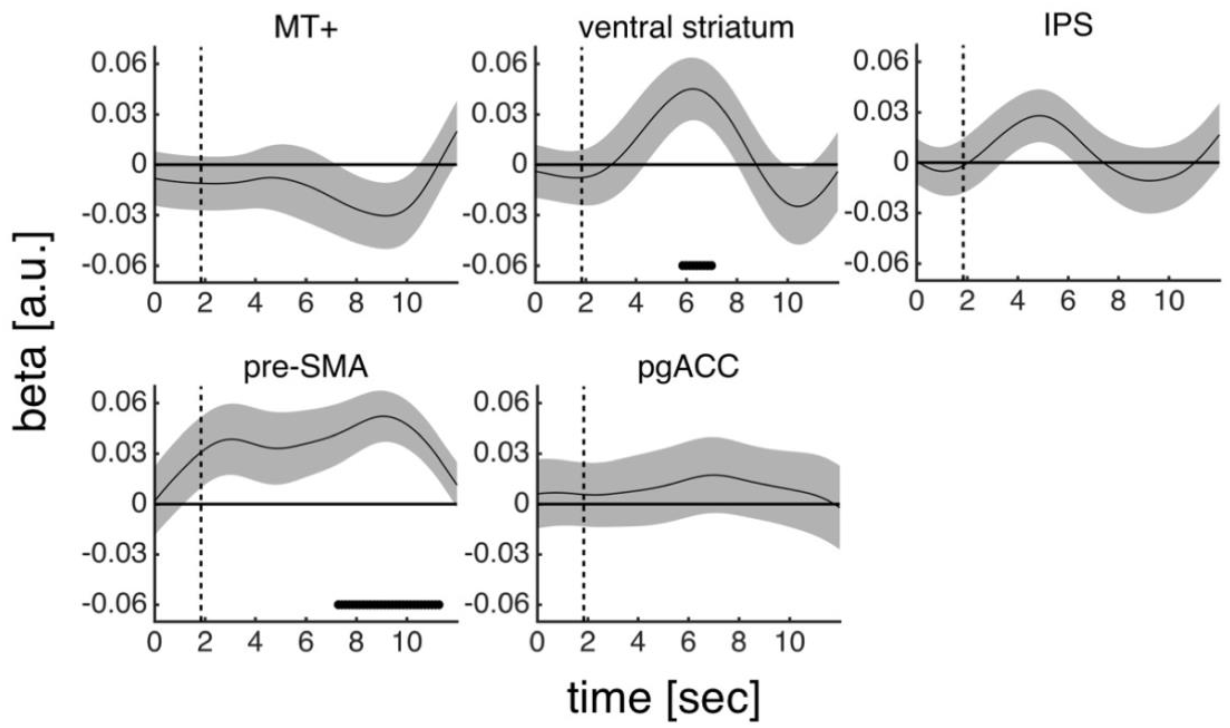
Encoding of reward in ROI activity time courses. General linear model analysis of the effects of reward magnitude (high versus low) on ROI activity time courses. Dots below time course indicate significant excursion of *t*-statistics assessed using two-tailed permutation tests. Vertical dashed line indicates the onset of the reward magnitude cue. Data, which only include confidence trials (15% of trials), are represented as group mean ± SEM.

**Figure S8.**
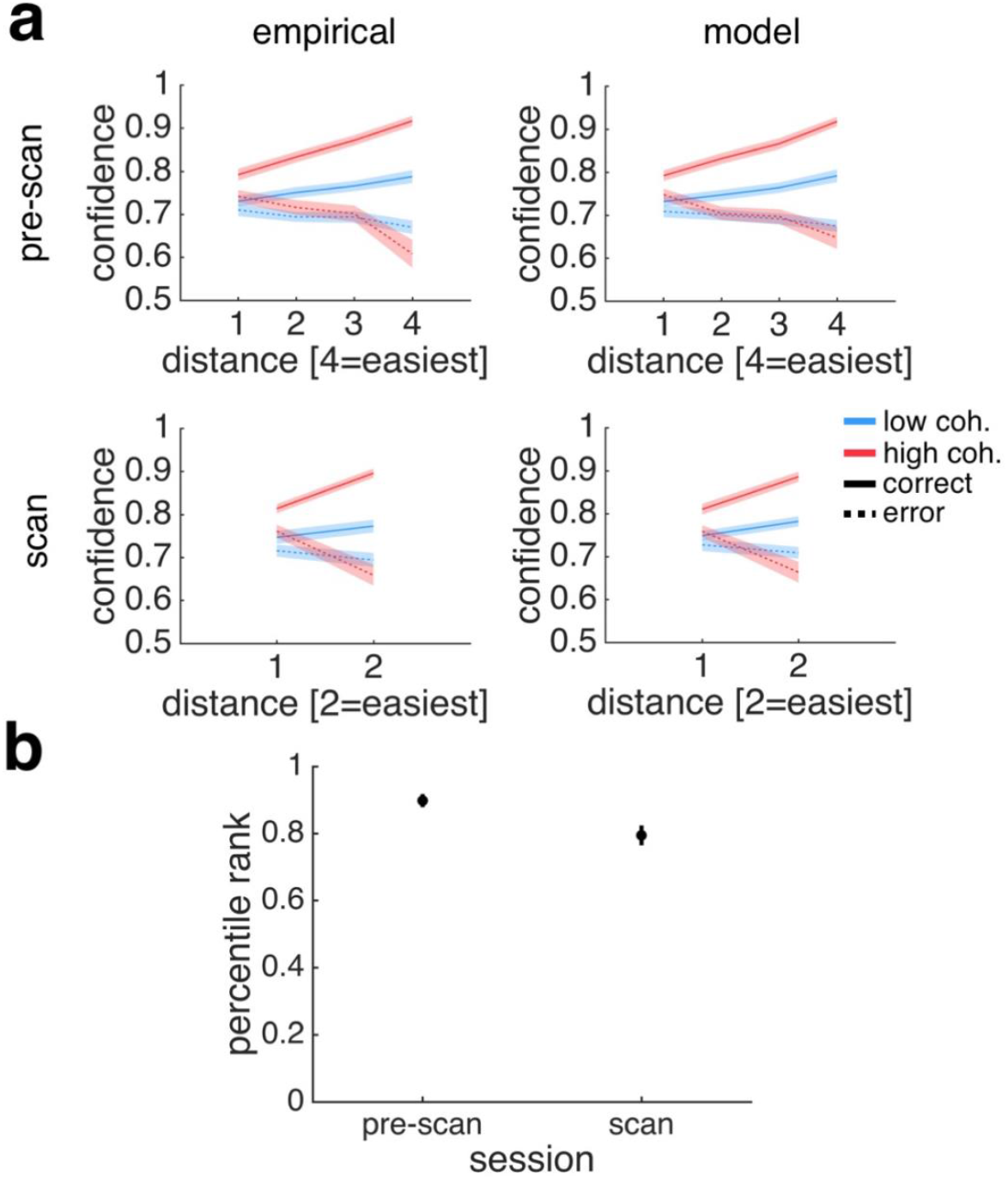
Evaluation of model of subjective confidence. **a**, Model qualitatively reproduces behaviour. The plots show *(left)* empirically observed and *(right)* model-derived confidence in the conditions of our factorial design split by choice accuracy (distance × coherence × choice accuracy) in *(top)* the pre-scan and *(bottom)* the scan session. The model was fitted to data from the pre-scan session and then used to create out-of-sample predictions about the scan session. We show empirical and predicted data from confidence trials only (15% of trials) for the scan session. **b**, Model provided good fits and generalised between sessions. For each subject, *s* = *i*, we computed the likelihood of the data under a model, *m*, fitted to the subject’s own pre-scan data, *m_s=i_*, and under the models fitted to the pre-scan data of every other subject, *m_s≠i_*. The plot shows (y-axis) the percentile rank of the likelihood under a subject’s own model, *m_s=I_*, compared to all other models, *m_s≠i_*, for (x-axis) the pre-scan and the scan sessions. Here higher rank indicates better model fits. As described in **Fig. 4a**, the model predicts, for each trial, the probability that a given response is made. Model predictions were evaluated by summing the likelihood of the observed trial-by-trial responses under this probability distribution. **a-b**, Data are represented as group mean ± SEM.

**Figure S9.**
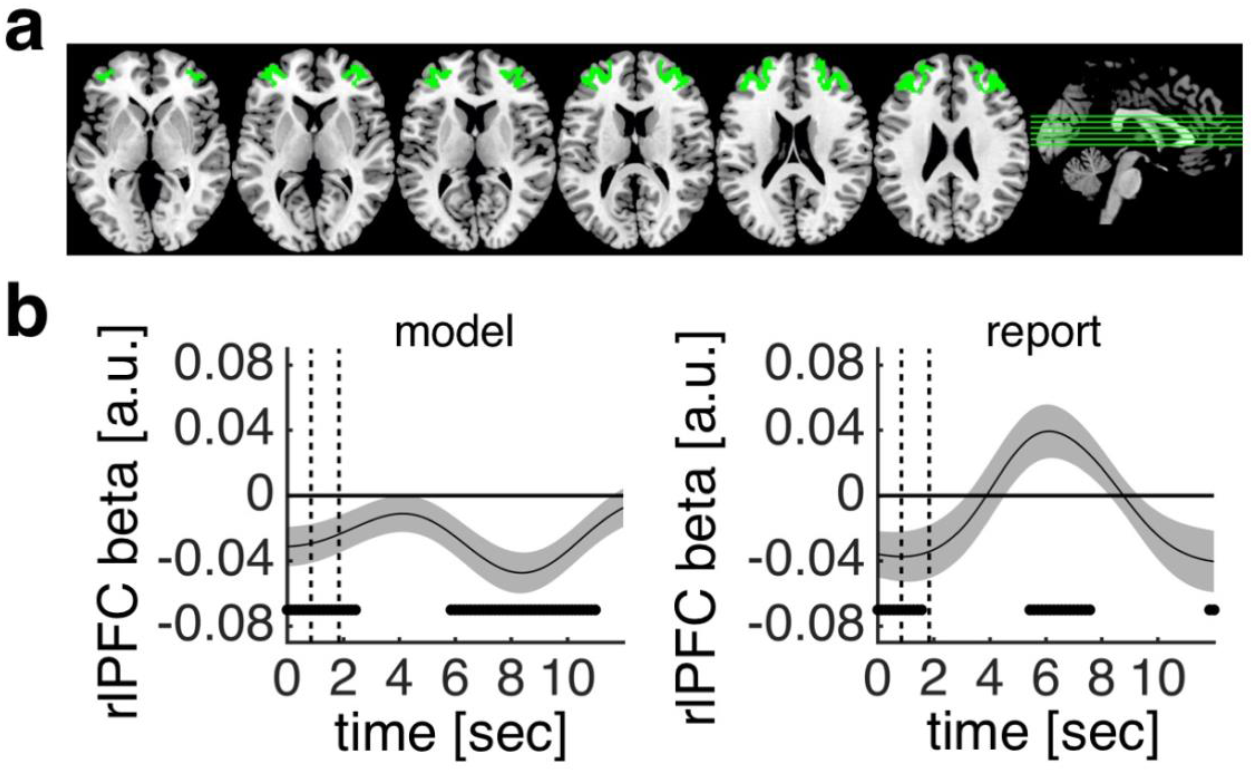
Activity in rlPFC predicts confidence estimates. **a**, ROI mask for rlPFC. **b**, General linear model analysis of encoding of model-derived subjective confidence (all trials) and reported confidence (confidence trials) in pgACC activity time courses. Dots below time course indicate significant excursion of *t*-statistics assessed using two-tailed permutation tests. Vertical dashed lines indicate the onsets of the motion stimulus and the choice phase. Data are represented as group mean ± SEM.

**Figure S10.**
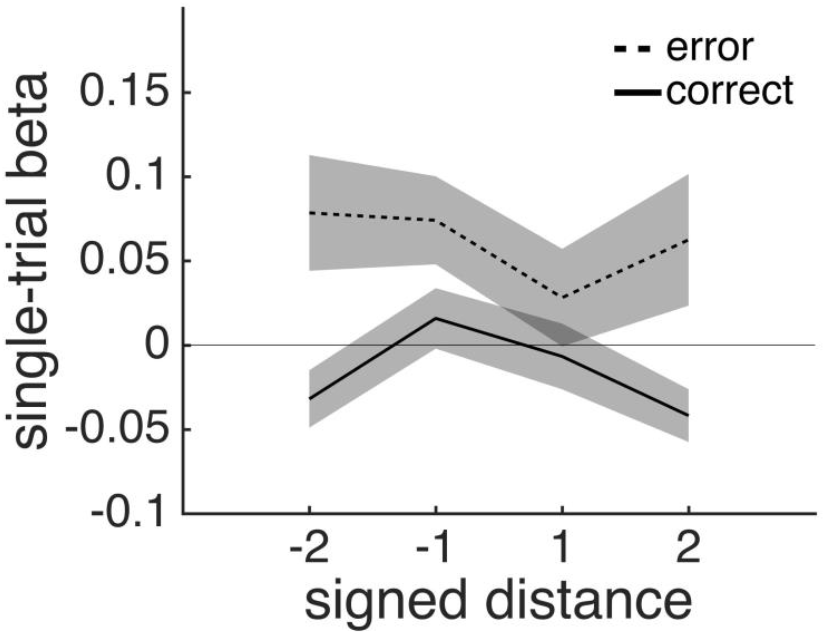
Activity in pre-SMA as a function of boundary distance and choice accuracy. Single-trial pre-SMA activity estimates shows an ‘X’-pattern as a function of signed distance (negative: CCW; positive: CW) and choice accuracy (dashed: error; solid: correct): activity is, on correct trials, lower for larger distances but, on error trials, higher for larger distances. This ‘X’-pattern is expected under a model which tracks the *inverse* of the absolute distance between sensory evidence and a choice boundary. Data are represented as group mean ± SEM.

**Figure S11.**
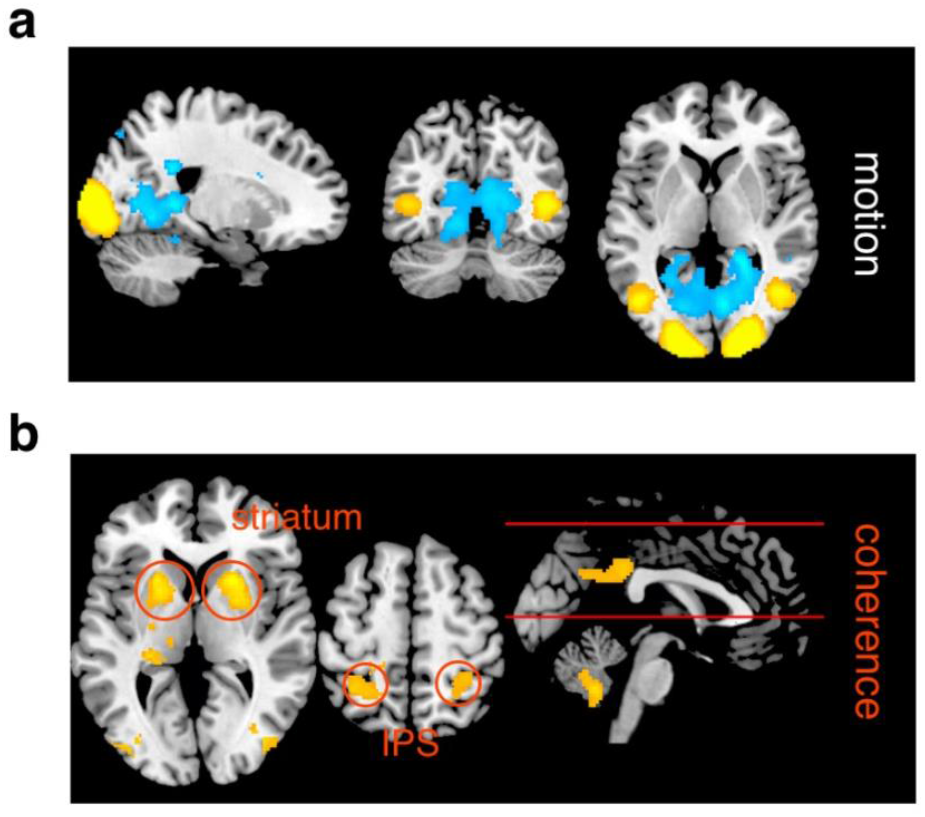
Neural responses to coherently moving dots. **a**, Whole-brain analysis of contrast between coherently moving and static dots in motion-localiser scan. **b**, Whole-brain analysis of main effect of coherence masked by coherent-motion contrast. We applied an exclusive mask constructed from the contrast shown in panel a (thresholded at p < .05, uncorrected) to GLM1. **a-b**, Cluster colours denote positive (warm) and negative (cold) effects. Clusters are significant at *p* < .05, FWE-corrected for multiple comparisons; cluster-defining threshold: *p* < .001, uncorrected. Images are shown at *p* < .001, uncorrected.

**Table S1.** Summary of significant activations for GLM1 post-masking. All reported activations are significant at *p* < .05, FWE-corrected for multiple comparisons; cluster-defining threshold: *p* < .001, uncorrected. FWE: familywise error. MNI: Montreal Neurological Institute. L: left. R: right. pre-SMA: pre-supplementary motor area. pgACC: perigenual anterior cingulate cortex.

| contrast | label | voxels at $p < .001$ | peak z-score | $p$ (cluster FWE corrected) | peak voxel MNI coordinates | | | |
| --- | --- | --- | --- | --- | --- | --- | --- | --- |
| high > low coh<br>*exclusive mask:*<br>*distance and interaction* | striatum | 1423 | 5.41 | <.001 | 26 | 10 | -6 | R |
|  | extrastriate (incl. MT+) | 2675 | 5.36 | <.001 | 28 | -84 | 8 | R |
|  | striatum | 1505 | 5.14 | <.001 | -24 | 10 | -10 | L |
|  | extrastriate (incl. MT+) | 1635 | 4.72 | <.001 | -40 | -78 | 2 | L |
|  | posterior parietal | 373 | 4.54 | .001 | 38 | -26 | 40 | R |
|  | cerebellum | 428 | 4.42 | <.001 | -4 | -56 | -42 | LR |
|  | posterior cingulate | 336 | 4.39 | .001 | 12 | -38 | 30 | LR |
|  | posterior parietal | 394 | 4.25 | .001 | -32 | -50 | 62 | L |
| high < low coh<br>*exclusive mask:*<br>*distance and interaction* | -----none----- |  |  |  |  |  |  |  |
| high > low dis<br>*exclusive mask:*<br>*coherence and interaction* | posterior cingulate | 214 | 4.91 | .014 | -12 | -16 | 48 | LR |
|  | early visual | 178 | 3.77 | .016 | -16 | -98 | 2 | L |
| high < low dis | pre-SMA | 169 | 4.34 | .020 | 4 | 20 | 48 | LR |
| interaction +<br>*inclusive mask:*<br>*coherence and distance* | pgACC | 932 | 4.82 | <.001 | -2 | 44 | 10 | LR |
| interaction –<br>*inclusive mask:*<br>*coherence and distance* | -----none----- |  |  |  |  |  |  |  |

**Table S2.**
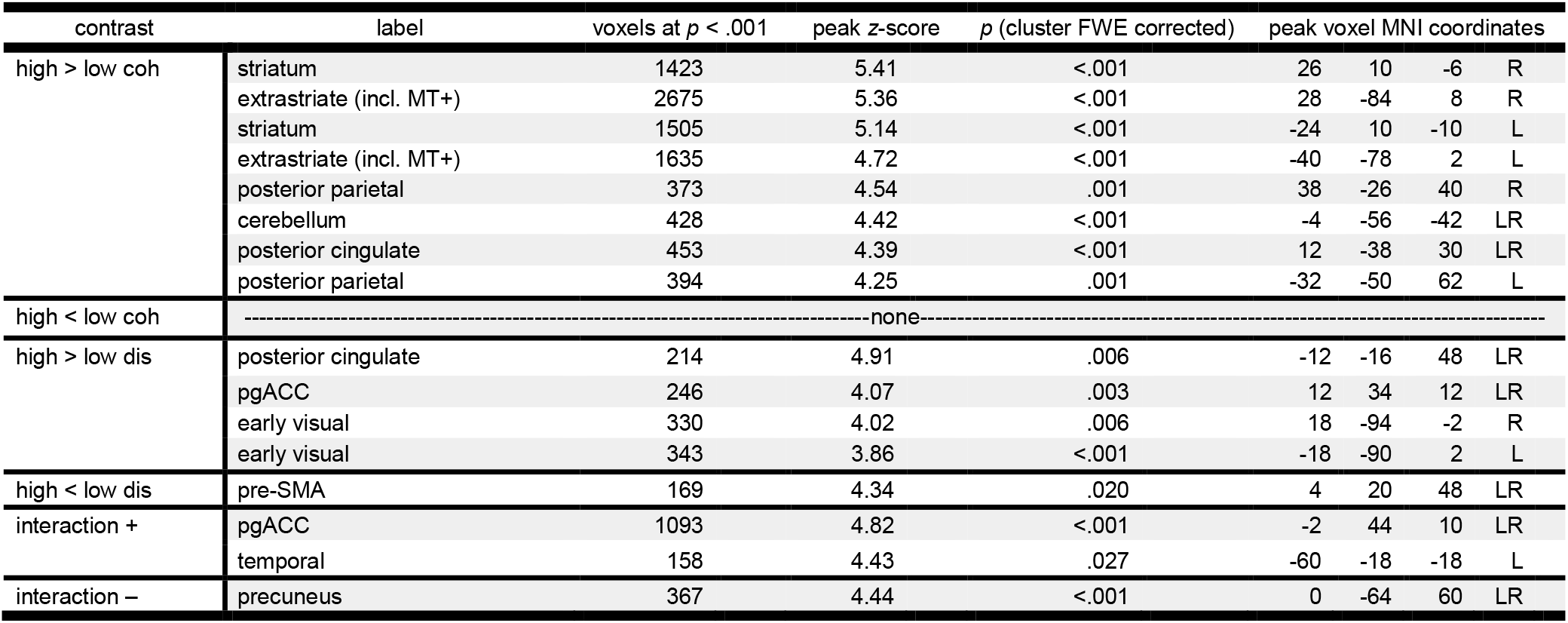
Summary of significant activations for GLM1 pre-masking. All reported activations are significant at *p* < .05, FWE-corrected for multiple comparisons; cluster-defining threshold: *p* < .001, uncorrected. FWE: familywise error. MNI: Montreal Neurological Institute. L: left. R: right. pgACC: perigenual anterior cingulate cortex. pre-SMA: pre-supplementary motor area.

**Table S3.** Summary of significant activations for GLM1C (correct trials only) post-masking. All reported activations are significant at *p* < .05, FWE-corrected for multiple comparisons; cluster-defining threshold: *p* < .001, uncorrected. FWE: familywise error. MNI: Montreal Neurological Institute. L: left. R: right. pgACC: perigenual anterior cingulate cortex. vmPFC: ventromedial prefrontal cortex.

| contrast | label | voxels at $p < .001$ | peak z-score | $p$ (cluster FWE corrected) | peak voxel MNI coordinates | | | |
| --- | --- | --- | --- | --- | --- | --- | --- | --- |
| high > low coh<br>*exclusive mask:*<br>*distance and interaction* | extrastriate (incl. MT+) | 791 | 4.89 | <.001 | 44 | -62 | 4 | R |
|  | striatum | 467 | 4.81 | <.001 | -22 | 12 | -10 | L |
|  | striatum | 610 | 4.77 | <.001 | 20 | 16 | -4 | R |
| high < low coh<br>*exclusive mask:*<br>*distance and interaction* | -----none----- |  |  |  |  |  |  |  |
| high > low dis<br>*exclusive mask:*<br>*coherence and interaction* | pgACC/vmPFC | 570 | 5.00 | .001 | 10 | 38 | 12 | LR |
|  | insula | 159 | 4.57 | .023 | -30 | 20 | -18 | L |
|  | insula | 140 | 4.39 | .040 | 32 | 14 | -10 | R |
|  | posterior cingulate | 150 | 4.38 | .030 | 0 | -12 | 40 | LR |
| high < low dis<br>*exclusive mask:*<br>*coherence and interaction* | -----none----- |  |  |  |  |  |  |  |
| interaction +<br>*inclusive mask:*<br>*coherence and distance* | pgACC | 252 | 4.28 | .002 | 4 | 44 | 12 | LR |
| interaction –<br>*inclusive mask:*<br>*coherence and distance* | -----none----- |  |  |  |  |  |  |  |

**Table S4.** Summary of significant activations for GLM1C (correct trials only) pre-masking. All reported activations are significant at *p* < .05, FWE-corrected for multiple comparisons; cluster-defining threshold: *p* < .001, uncorrected. FWE: familywise error. MNI: Montreal Neurological Institute. L: left. R: right. pgACC: perigenual anterior cingulate cortex.

| contrast | label | voxels at $p < .001$ | peak z-score | $p$ (cluster FWE corrected) | peak voxel MNI coordinates | | | |
| --- | --- | --- | --- | --- | --- | --- | --- | --- |
| high > low coh | extrastriate (incl. MT+) | 2050 | 4.89 | <.001 | 44 | -62 | 4 | R |
|  | striatum | 618 | 4.81 | <.001 | -22 | 12 | -10 | L |
|  | striatum | 768 | 4.77 | <.001 | 20 | 16 | -4 | R |
|  | extrastriate (incl. MT+) | 1307 | 4.23 | <.001 | -24 | -88 | 20 | L |
|  | posterior parietal | 278 | 4.01 | .002 | -24 | -52 | 58 | L |
|  | posterior parietal | 172 | 3.97 | .027 | 30 | -48 | 62 | R |
| high < low coh | -----none----- |  |  |  |  |  |  |  |
| high > low dis | pgACC | 610 | 5.00 | <.001 | 10 | 38 | 12 | LR |
|  | insula | 229 | 4.57 | .004 | -30 | 20 | -18 | L |
|  | insula | 182 | 4.41 | .012 | 32 | 14 | -10 | R |
|  | posterior cingulate | 293 | 4.38 | <.001 | 0 | -12 | 40 | LR |
|  | premotor | 220 | 4.36 | .004 | -46 | -18 | 52 | L |
| high < low dis | -----none----- |  |  |  |  |  |  |  |
| interaction + | pgACC | 309 | 4.28 | <.001 | 4 | 44 | 12 | LR |
| interaction - | precuneus | 520 | 4.41 | <.001 | 10 | -66 | 64 | L |

